# Boosting alignment accuracy through adaptive local realignment

**DOI:** 10.1101/063131

**Authors:** Dan DeBlasio, John Kececioglu

## Abstract

**Motivation:** While mutation rates can vary across the residues of a protein, when computing alignments of protein sequences the same setting of values for substitution score and gap penalty parameters is typically used across their entire length. We provide for the first time a new method called *adaptive local realignment* that automatically uses diverse parameter settings in different regions of the input sequences when computing multiple sequence alignments. This allows parameter settings to adapt to more closely match the local mutation rate across a protein.

**Method:** Our method builds on our prior work on global alignment *parameter advising* with the `Facet` alignment accuracy estimator. Given a computed alignment, in each region that has low estimated accuracy, a collection of candidate realignments is generated using a precomputed set of alternate parameter settings. If one of these alternate realignments has higher estimated accuracy than the original subalignment, the region is replaced with the new realignment, and the concatenation of these realigned regions forms the final alignment that is output.

**Results:** Adaptive local realignment significantly improves the quality of alignments over using the single best default parameter setting. In particular, this new method of *local advising*, when combined with prior methods for *global advising*, boosts alignment accuracy by as much as 26% over the best default setting on hard-to-align benchmarks (and by 6.4% over using global advising alone).

**Availability:** A new version of the `Opal` multiple sequence aligner that incorporates adaptive local realignment using Facet for parameter advising, is available free for non-commercial use at http://facet.cs.arizona.edu.

## 1 Introduction

Since the 1960s it has been known that proteins can have distinct mutation rates at different locations along the molecule (Fitch and Margoliash, 1967). The amino acids at some positions in a protein may stay unmutated for long periods of time, while other regions change a great deal (often called “hypermutable” regions). This has led to methods in phylogeny construction that take variable mutation rates into account when building trees from sequences (Yang, 1993). In multiple sequence alignment, however, to our knowledge variation in mutation rates across sequences has yet to be exploited to improve alignment accuracy. Multiple sequence alignments are typically computed using a single setting of values for the parameters of the alignment scoring function. This single parameter setting affects how residues across a protein are aligned, and implicitly assumes a uniform mutation rate. In contrast, the approach of this paper identifies alignment regions that may be misaligned under a single parameter setting, and finds alternate parameter settings that may more closely match the local mutation rate of the sequences.

**Figure 1.**
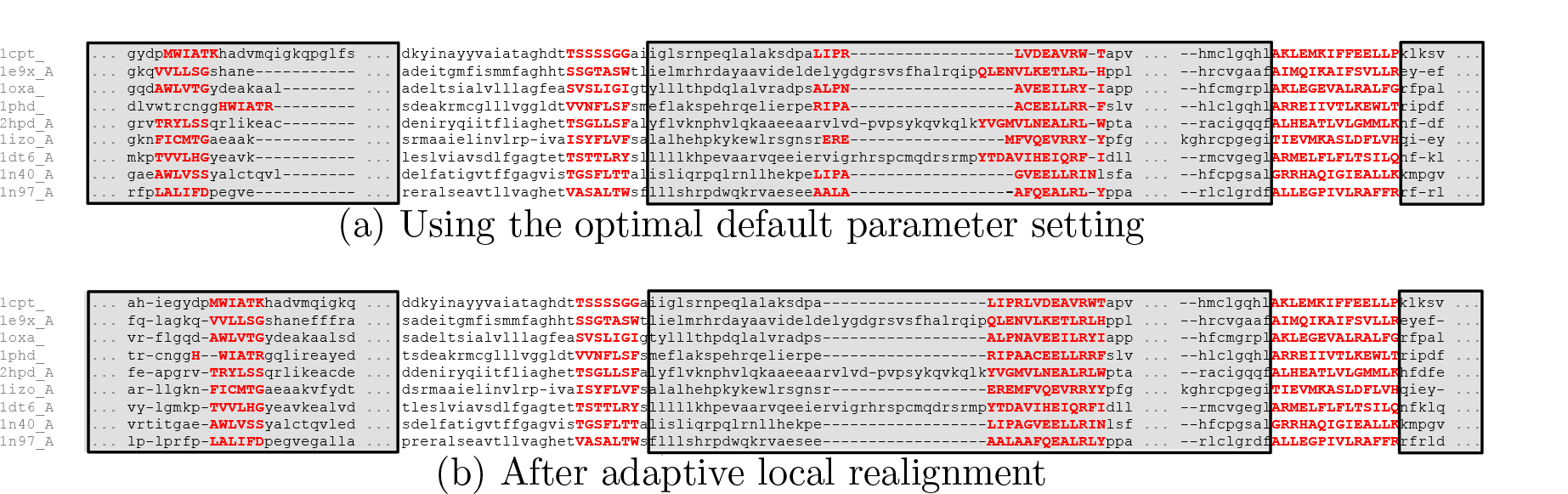
Impact of adaptive local realignment. The figure shows portions of an alignment of benchmark `BB11007` from the `BAliBASE` suite, where the highlighted amino acids in red uppercase are from the core columns of the reference alignment, which should be aligned in a correct alignment, (a) The alignment computed by Opal using its optimal default parameter setting (`VTML200`, 45,11, 42,40) across the sequences, with an accuracy of 89.6%. The regions of the alignment in gray boxes are automatically selected for realignment, (b) The outcome of using adaptive local realignment, with an improved accuracy of 99.6%. The realignments of the three regions use alternate parameter settings (`BL0SUM62`, 45, 2,45, 42), (Figure 1`BL0SUM62`, 95, 38,40,40), and (`VTML200`,45,18,45,45), respectively, which increase the accuracy of these regions.

We present a method that takes a given alignment and attempts to improve its overall accuracy by replacing sections of it with better subalignments, as demonstrated in Figure 1. The top alignment of the figure was computed using a single parameter setting: the optimal default setting of the `Opal` aligner (Wheeler and Kececioglu, 2007). The bottom alignment is obtained by our new method, taking the top alignment, automatically identifing the sections in gray boxes, and realigning them using alternate parameter settings, as described later in Section 3. This increases the overall alignment accuracy by 10%, as most of the misaligned core blocks (highlighted in red uppercase) are now corrected.

### Related work

Methods that partition a set of sequences to align or realign them can be divided in two categories, based on the type of partition. *Vertical* realigners cut the input sequences into substrings, and once these shorter substrings are realigned, they stitch their alignments together. *Horizontal* realigners split an alignment into groups of whole sequences, which are then merged together by realigning between groups, possibly using each group’s induced subalignment.

`Crumble` and `Prune` (Roskin *et al.*, 2011) is a pair of algorithms for performing both vertical (`Crumble`) and horizontal (`Prune`) splits on an input set of sequences. During the `Crumble` stage, a set of constraints is found that anchor the input sequences together, and the substrings or blocks between these anchor points are aligned. Once the disjoint blocks of the sequences are aligned, they are then fused by aligning their overlapping anchor regions. During the `Prune` stage, smaller groups of sequences are aligned that correspond to subtrees of the progressive aligner’s guide tree. The subset of sequences in a subtree is then replaced by their alignment’s consensus sequence in the remaining steps of progressive alignment. The original subalignments of the groups are finally reinserted to form the output alignment. Replacing a group of sequences by their consensus sequence during alignment reduces the number of sequences that are aligned at any one time. The objective for splitting sequences both vertically and horizontally within `Crumble` and `Prune` is to reduce time and space usage to make feasible the alignment of large numbers of long sequences, rather than to improve alignment accuracy.

`ReAligner` (Anson and Myers, 1997) is a horizontal realignment method for improving DNA sequence assembly by removing and then realigning sequencing reads. If a read is initially misaligned in the assembly it may be corrected when the read is removed and realigned.This realignment process is repeated over all reads to continually refine the assembly.

Gotoh (1993) presented several horizontal methods for heuristically aligning two multiple sequence alignments, which he called “group-to-group” alignment. This could be used for alignment construction in a progressive aligner, proceeding bottom-up over the guide tree and applying group-to-group alignment at each node, or for polishing an existing alignment by assigning sequences to two groups and using it to realign the groups.

`AlignAlign` (Kececioglu and Starrett, 2004) is a horizontal method that implements an exact algorithm for optimally aligning two multiple sequence alignments under the sum-of-pairs scoring function with affine gap costs. This optimal group-to-group alignment algorithm, used for both alignment construction and alignment polishing, forms the basis of the `Opal` aligner (Wheeler and Kececioglu, 2007).

By comparison, adaptive local realignment is a vertical approach that aims to improve alignment accuracy, applies to any alignment method that has tunable parameters, and to our knowledge is the first approach to alignment that can automatically adapt to varying mutation rates along a protein.

### Plan of the paper

The next section provides the necessary background on parameter advising, which is basic to our adaptive local realignment technique. A parameter advisor selects a parameter setting for an aligner from a small set of choices drawn from a larger universe of all possible settings, using an alignment accuracy estimator to select its choice. Section 3 then describes our adaptive local realignment method, which can be viewed as a form of local parameter advising, and discusses how it interacts with global parameter advising. Section 4 experimentally evaluates our approach, and compares it with prior methods for advising. Finally, Section 5 gives conclusions and offers directions for further research.

## 2 Background on parameter advising

To make the paper self-contained, we briefly review our prior work on how to learn an alignment advisor. We first review the concept of *parameter advising*, which requires an estimator of alignment accuracy and a set of parameter choices for the advisor, and then summarize our prior techniques for learning both an estimator and an advisor set.

Note that while we present here on how to find advisor estimators and advisor sets based on training data for the user the advisor estimator and sets are precomputed with no need for retraining.

### 2.1 Global parameter advising

The goal of parameter advising is to find the parameter setting for an aligner that yields the most accurate alignment of a given set of input sequences. The accuracy of a computed alignment is measured with respect to the “correct” alignment of the sequences (which often is not known). For special benchmark sets of protein sequences, the gold-standard alignment of the proteins, called their *reference alignment*, is usually obtained through structural alignment by finding the best superposition of the known three-dimensional structures of the proteins. Columns of the reference alignment that contain a residue from every protein in the set (where a *residue* is the amino acid at a particular position in a protein), and for which the residues in the column are all mutually close in space in the superposition of the structures, are called *core columns*. Runs of consecutive core columns are called *core blocks*, which represent the regions of the structural alignment with the highest confidence of being correct. Given such a reference alignment with identified core blocks, the *accuracy* of a different, computed alignment is the fraction of the pairs of residues aligned in the core blocks of the reference alignment that are also aligned in the computed alignment. (So a computed alignment of 100% accuracy completely agrees with the reference on its core blocks, though it may disagree elsewhere.) The best computed alignment is one of highest accuracy, and the task of a parameter advisor is to find a setting of the tunable parameters of an aligner that yields an accurate output alignment. This setting can be highly input dependent, as the best choice of parameter values for an aligner can vary for different sets of input sequences.

When aligning sequences in practice, a reference alignment is almost never known, in which case the true accuracy of a computed alignment cannot be measured. Instead our parameter advisors rely on an accuracy *estimator E* that for an alignment *A*, gives a value *E*(*A*) in the range [0,1] that estimates the true accuracy of alignment *A*. An estimator should be efficiently computable and positively correlated with true accuracy.

To choose a parameter setting, an advisor takes a set of choices *P*, where each *parameter choice p* ∈ *P* is a vector that assigns values to all the tunable parameters of an aligner, and picks the choice that yields a computed alignment of highest estimated accuracy.

Formally, given an accuracy estimator *E* and a set *P* of parameter choices, a *parameter advisor* tries each parameter choice *p* ∈ *P*, invokes an aligner to compute an alignment *A_p_* using choice *p*, and then selects the parameter choice *p*\* that has maximum estimated accuracy: *p*\* ∈ argmax_*p* ∈ *P*_ {*E*(*A_p_*)}. Since the advisor runs the aligner |*P*| times on a given set of input sequences, a crucial aspect of parameter advising is finding a small set *P* for which the true accuracy of the output alignment *A_p*_* is high.

To construct a good advisor, we need to find a good estimator *E* and a good set *P*. The estimator and advisor set are learned on training data consisting of benchmark sets of protein sequences for which a reference alignment is known. The learning procedure tries to find an estimator *E* and set *P* that maximize the true accuracy of the resulting advisor on this training data, which we subsequently assess on separate testing data.

Note that the process of advising is fast: for a set *P* of *k* parameter choices, advising involves computing *k* alignments under these choices, which can be done in parallel, evaluating the estimator on these *k* alignments, and taking a max. The separate process of training an advisor, by learning an estimator and an advisor set as we review next, is done once, off-line, before any advising takes place.

### 2.2 Learning an accuracy estimator

Kececioglu and DeBlasio (DeBlasio *et al.*, 2012; Kececioglu and DeBlasio, 2013) present an efficient approach for learning an accuracy estimator that is a linear combination of real-valued alignment feature functions, based on solving a large-scale linear programming problem. This approach resulted in the `Facet` estimator (DeBlasio and Kececioglu, 2014a), which is currently the most accurate estimator for parameter advising (Kececioglu and DeBlasio, 2013; DeBlasio and Kececioglu, 2015b).

This approach assumes we have a collection of *d* real-valued feature functions *g*_1_(*A*),…, *g_d_*(*A*) on alignments *A*, where these functions *g_i_* are positively correlated with true accuracy. The alignment accuracy estimator *E* is a linear combination of these functions, *E*(*A*) = ∑_1≤*i*≤*d*_ *c_i_ g_i_*(*A*), where the coefficents *c_i_* specify the estimator *E*. When the feature functions have range [0,1] and the coefficients form a convex combination, the resulting estimator *E* will also have range [0,1]. `Facet` uses a collection of five feature functions, many of which make use of predicted secondary structure for the protein sequences (Kececioglu and DeBlasio, 2013). Figure 3 shows the relationship to true accuracy of both the `Facet` and `TCS` (Chang *et al.*, 2014) estimators. To be able to distinguish good from bad alignments an estimator should have a steep slope and very little spread. While the `TCS` estimator has higher slope it has quite a bit of spread. In contrast the `Facet` estimator has much less spread but a less steep slope, and we have found that this is more effective in ranking alignments for parameter advising.

The features we use in `Facet` are a mixture of canonical measure of alignment quality such as Amino Acid Identity and novel non-local features of alignment that correlate with true accuracy. Many of the most accurate features use protein secondary structure, these include secondary structure identity and Secondary Structure Blockiness. Blockiness finds a packing of contiguous groups of amino acids with the same structure label. The other feature functions used in the `Facet` estimator are: Secondary Structure Identity, Secondary Structure Agreement, Gap Open Density, and Core Column Percentage, A full description of the features can be found in Kececioglu and DeBlasio (2013).

**Figure 2.**
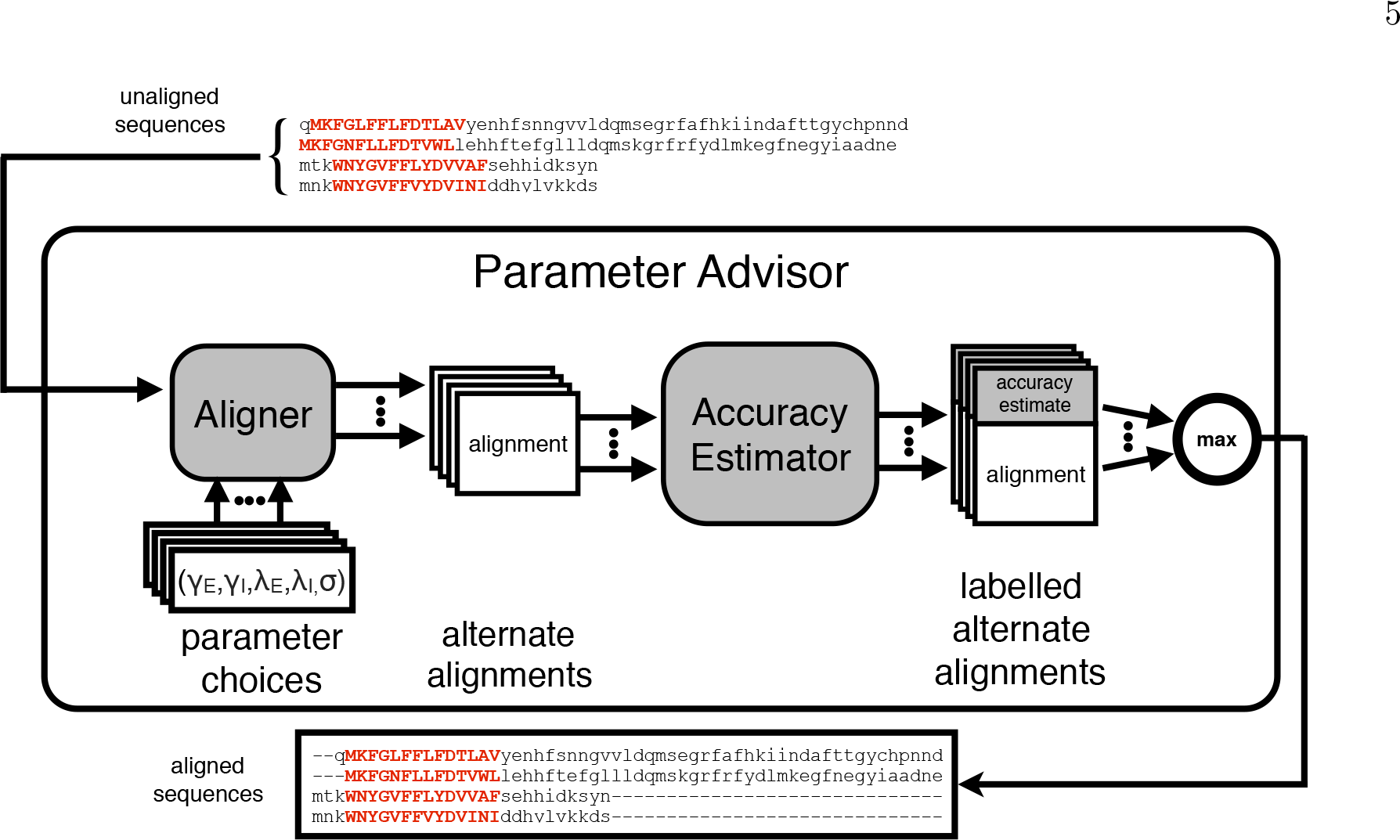
The parameter advising process. For an input set of sequences, a parameter advisor first invokes the aligner for each of a collection of independent parameter choices. Each parameter choice when used with the aligner produces an alternate alignment of the sequences. An accuracy estimator is then used to label each of the alternate alignments with an accuracy estimate. The advisor then returns the alternate alignment with the highest accuracy estimate.

A parameter advisor uses the estimator to effectively rank alignments, so an estimator just needs to be monotonic in true accuracy. The *difference-fitting* approach learns the coefficients of an estimator that is close to monotonic by fitting the estimator to differences in true accuracy for pairs of training alignments.

Let function *F*(*A*) give the *true accuracy* of alignment *A*, and set 𝒫 be a collection of ordered pairs of alignments from training data, where every pair (*A*,*B*) ∈ 𝒫 satsifies *F*(*A*) < *F*(*B*). Difference fitting tries to find an estimator *E* that increases at least as much as accuracy *F* on the pairs in 𝒫, by minimizing the amount that *E* falls short. Formally, we find the estimator *E*\* given by the vector of coefficients 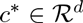 that minimizes 
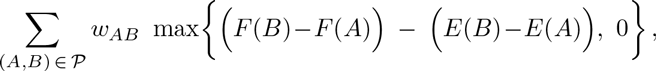
where *w_AB_* weights the above error for a pair (*A*, *B*). Finding the optimal coefficients *c*\* can be reduced to solving a linear programming problem as follows.

The linear program has a variable *c_i_* for each estimator coefficient, and an error variable *e_AB_* for each pair (*A, B*) ∈ 𝒫. The constraints are *c_i_* ≥ 0 and ∑_*i*_ *c_i_* = 1, which ensure the coefficients form a convex combination, together with *e_AB_* ≥ 0, and 
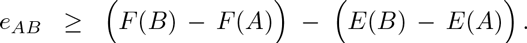
(Note that the expressions *E*(*A*) and *E*(*B*) are linear in the variables *c_i_*, while the quantities *F*(*A*) and *F*(*B*) are constants.) The linear program then minimizes the objective function ∑_(*A*,*B*)∈𝒫_ *w_AB_ e_AB_*.

**Figure 3.**
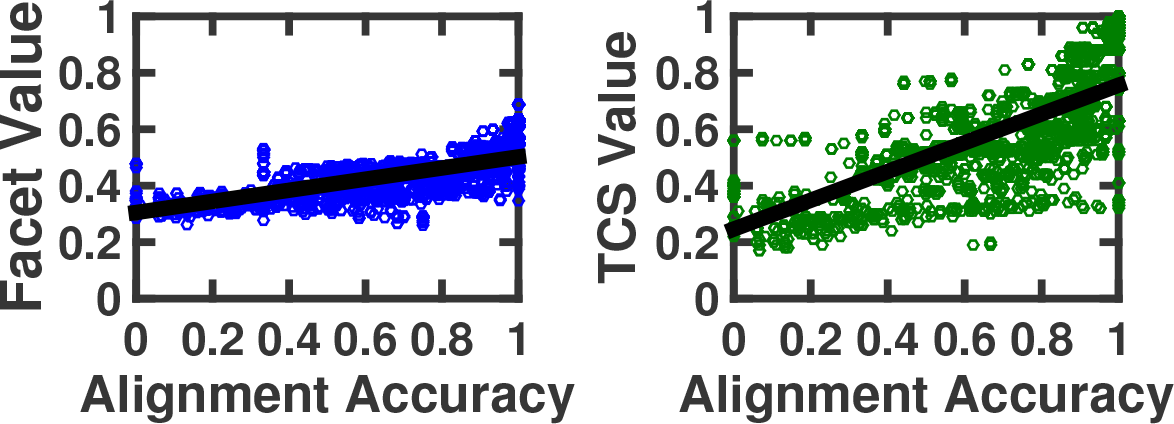
*Relationship of estimators to true accuracy*. Each point in a scatterplot corresponds to an alignment whose true accuracy is on the horizontal axis, and whose value under a given estimator is on the vertical axis. Both scatterplots show the same set of 3,000 alignments under the accuracy estimators `Facet` (Kececioglu and DeBlasio, 2013) and `TCS` (Chang *et al.*, 2014).

For the linear program to be of manageable size for a large number of training alignments, the set 𝒫 of pairs must be quite sparse. Kececioglu and DeBlasio (2013) describe how to find a good sparse set 𝒫 together with a good set of weights *w_A B_* by an efficient graph algorithm.

Learning an accuracy estimator with *d* feature functions using a set 𝒫 of *p* pairs, involves solving the above linear program with 𝒫 + *d* variables and Θ(*p* + *d*) inequalities. Evaluating the `Facet` estimator on an alignment with *m* sequences and *n* columns, after secondary structure has been predicted for the protein sequences, takes Θ(*m*^2^*n*) time.

### 2.3 Learning an advisor set

DeBlasio and Kececioglu (2014b, 2015b) present an efficient approximation algorithm for learning a near-optimal set of parameter choices for an advisor that uses a given estimator. The approximation algorithm follows a greedy strategy, so we call the sets found by the approximation algorithm *greedy sets*, in contrast to *exact sets* that are optimal for the training data, and which can be found by exhaustive search for small instances. These greedy sets tend to generalize better than exact sets, with the remarkable behavior that the greedy sets often outperform exact sets on testing data DeBlasio and Kececioglu (2015b).

The problem of learning an optimal set *P* of parameter choices for an advisor is formulated as follows. Let *U* be the *universe* of possible parameter choices that might be included in advisor set *P*. The training data is a collection of *reference alignments R_i_*, one for each benchmark *B_i_*, and a collection of *alternate alignments A_ij_*, where each alignment *A_ij_* is computed by running the aligner on the sequences in benchmark *B_i_* using parameter choice *j* ∈ *U*. By comparing each alternate alignment to the reference alignment for its benchmark, we can measure the *true accuracy a_ij_* of each alignment *A_ij_*. Details of the parameter universe used for the `Opal` aligner can be found in Section 4.

For a candidate set *P* ⊆ *U* of parameter choices for an advisor that uses estimator *E*, the set of parameter choices from *P* that could potentially be *output* by the advisor on benchmark *B_i_* is *𝒪_i_* (*P*) = argmax_*j* ∈ *P*_ {*E* (*A_ij_*)}, where the argmax gives the set of parameter choices *j* ∈ *P* that are tied for maximizing the estimator *E* on the benchmark. The advisor could output any parameter choice from *𝒪_i_*(*P*), as all of them appear equally good under the estimator, so we assume the advisor selects a choice uniformly at random from this set. Then the *expected accuracy* achieved by the advisor on benchmark *B_i_* using parameter set *P* is 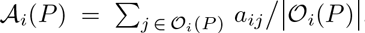, where again *a_ij_* is the true accuracy of alignment *A_ij_*.

In learning an advisor set *P*, we seek a set *P* that maximizes the advisor’s expected accuracy 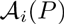 on the training benchmarks *B_i_*. Formally, we want a set P that maximizes the objective function 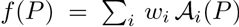, where *i* indexes the benchmarks, and *w_i_* is the weight placed on benchmark *B_i_*. (The benchmark weights correct for sampling bias in the training data, as discussed in Section 4.) In words, we want to find an advisor set *P* ⊆ *U* that maximizes the expected accuracy of the parameter choices selected by the advisor, averaged across weighted training benchmarks.

DeBlasio and Kececioglu (2015b) prove that for a given bound *k* on the size of advisor set *P*, finding an optimal set *P* ⊆ *U* with |*P*| ≤ *k* that maximizes objective *f* (*P*) is NP-complete. They further develop an efficient greedy approximation algorithm that is guaranteed to find near-optimal advisor sets, though for adaptive local realignment these *greedy sets* do not perform as well as the oracle sets we describe next.

#### 2.3.1 Oracle sets

A useful notion in parameter advising is the concept of an oracle (Wheeler and Kececioglu, 2007), which is a perfect advisor that has access to the true accuracy of an alignment. For a given advisor set *P*, an *oracle* selects parameter choice argmax_*p*∈*P*_{*F*(*A_p_*)}, where again function *F* gives the true accuracy of an alignment. (Equivalently, an oracle is an advisor that uses the perfect estimator *F*.) An oracle always picks the parameter choice that yields the highest accuracy alignment.

While an oracle is impossible to construct in practice, it gives a theoretical limit on the accuracy achievable by advising with a given set. Furthermore, if we can find the optimal advisor set for an oracle for a given cardinality bound *k*, which we call an *oracle set*, then the performance of an oracle on an oracle set gives a theoretical limit on how well advising can perform for a given bound *k* on the number of parameter choices.

Kececioglu and DeBlasio (2013) show that finding an oracle set is NP-complete, and give the following integer linear program for finding an optimal oracle set. In our prior notation, an optimal oracle set for cardinality bound *k* is a set *P* ⊆ *U* with |*P*| ≤ *k* that maximizes ∑_*i*_ *w_i_* max_*j*∈*P*_ *a_ij_*. To formulate this as an integer linear programming problem, let *y_j_* for all *j* ∈ *U*, and *x_ij_* for all benchmarks *B_i_* and all *j* ∈ *U*, be integer variables that either have the value 0 or 1, where *y_j_* = 1 iff parameter choice *j* is selected for oracle set *P*, and *x_ij_* = 1 iff choice *j* is used by the oracle on benchmark *B_i_*. The constraints are 0 ≤ *x_ij_* ≤ *y_j_* ≤ 1, ∑_*j*_ *y_j_* ≤ *k*, and for each benchmark *B_i_* the constraint ∑_*j*_ *x_ij_* = 1. The objective function is to maximize ∑_*i*_ *w*i** ∑_*j*_ *x_ij_ a_ij_*. An optimal solution to this integer linear program gives an optimal oracle set *P*\* = {*j* ∈ *U*: *y_j_* = 1}.

Learning an optimal oracle set of cardinality *k*, for a universe of *u* parameter choices and a training set of *t* benchmarks, involves solving the above integer linear program with Θ(*ut*) variables and Θ(*ut*) inequalities. Kececioglu and DeBlasio (2013) show that using the `CPLEX` integer linear programming solver, this formulation permits finding optimal oracle sets in practice even for cardinalities up to *k* = 25.

This previous work has focused on using parameter advising to choose the parameter setting for an entire alignment, which we call here *global parameter advising*. The next section presents our adaptive local realignment method, which leverages the ideas reviewed here to achieve what can be viewed as *local parameter advising*.

## 3 Adaptive local realignment

To overcome the issue of protein sequences being non-homogeneous and having regions that may require different alignment parameters we have developed a method we call *adaptive local realignment*. Adaptive local realignment uses some of the the same basic principles that have been shown to work well for global parameter advising. We apply the techniques described above locally to choose the best alignment parameters for a subset of columns of an alignment.

The adaptive local realignment method for an alignment can be broken down into two steps: (1) choosing regions of the alignment that are correctly aligned which we should save, and (2) producing a new alignment for those regions that are not correctly aligned.

Similar to global parameter advising local realignment relies on a set of alternate parameter choices and the accuracy estimator.

### 3.1 Identifying local realignment regions

Just as with global alignments we do not have a known reference alignment so we cannot simply identify the alignment columns that are recovered correctly in an input computed alignment. Therefore we are forced to use an accuracy estimator *E* to define the regions of a given alignment that are going to be saved (and which will be realigned). We calculate the estimated accuracy of a sliding window across the alignment (see Figure 4a). The window size is a fraction *w* ≤ 1 of the total length of the alignment. The window size *w* must be chosen carefully because the accuracy estimator has features that are global calculations of an alignment. A larger sliding window will provide more context at each position and should provide a better estimate of accuracy. At the same time, if the window is too large there will not be enough granularity to identify the separation points between correct and incorrectly aligned columns. To account for very short and very long alignments a minimum *w_min_* and maximum *w_max_* window size is specified.

We now have the scores for approximately 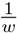 windows that overlap each column of an alignment. We calculate a score for each column as a sum of these scores weighted proportionally to the distance to the center column of that window (see Figure 4b). We use a gamma decay distribution with a decay factor *d* ≤ 1 centered on the middle column to weight the contribution of each window. As *d* approaches 1 a column gets equal weight from all covering windows. Conversely as it approaches 0 the score is dependent only on the window centered at that column.

We then calculate two thresholds *τ_B_* and *τ_S_* based on the percentage of columns from the original alignment we would like to keep *T_B_* and the percentage of columns we will use to seed realignment regions *T_S_*. The thresholds are set so that at least [*ℓT_B_*] columns have score that are above *τ_B_*, and least [*ℓT_S_*] columns have scores below *τ_S_*. All columns with scores ≥ *τ_B_* are labeled “barriers” and all columns with scores ≤ *τ_s_* are labeled as “seeds” (see Figure 4b).

To find alignment regions that will be realigned we start a region by including a seed column. This region is then extended to include any other seed or unlabeled column to the left and right. This expansion continues until a barrier column (or the end of the alignment) is reached in both directions.

The barrier columns will never be included in an alignment region that will be realigned. In this way we guarantee that at least *T_B_* percent of the columns from the original alignment will remain, and there will always be at least one alignment region to realign.

### 3.2 Local parameter advising on a region

During local advising we will construct a new alignment that contains all of the columns that are surrounded by only barrier columns and more accurate alignments of the columns covered by alignment regions (if a more accurate alignment can be found).

For each alignment region we extract the sub-alignment in the contained columns and calculate it’s `Facet` score (see Figure 4d). We will later compare other alternate alignments with this base accuracy. The unaligned subsequences of this region are then collected. This set of unaligned sequences becomes the input to parameter advising.

We use same parameter advising method described in Section 2.1 and Figure 2 with one exception. The `Opal` aligner has 5 tunable parameters: the replacement matrix as well as two internal and two terminal gap costs (we describe them in detail in Section 4). For those regions that do include the alignment terminals (the first or last column of the input alignment), the terminal gap penalties are replaced with the corresponding internal gap cost. For those regions that do include terminals we use the terminal gap penalty only on the one side that is terminal in the context of the global alignment. Note that an alignment region as we have defined it will never include both terminals.

**Figure 4.**
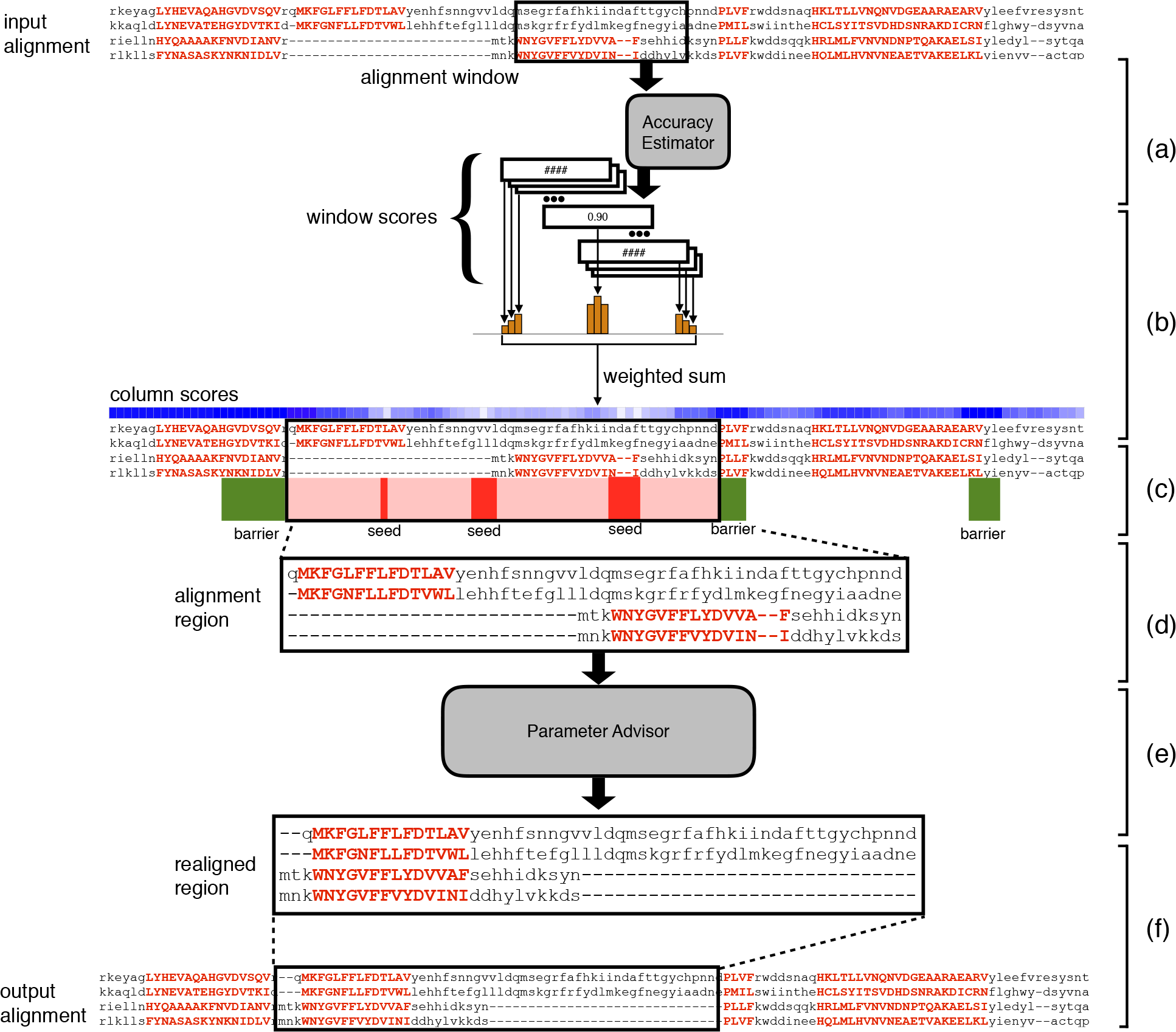
The adaptive local realignment process. (a) We calculate a `Facet` score for a sliding window across at the input alignment. (b) To calculate a score for each column from the set of window scores we use a weighted sum of the values for all windows that overlap that column. (c) Columns that a column score value greater than *τ_G_* are labeled as barriers and then columns with value less than *τ_B_* are used as seeds for realignment regions. (d) These seeds are then extended in both directions until they reach a barrier column to define a realignment region that is extracted from the alignment. (e) The unaligned subsequences defined by this region are then realigned using a parameter advisor. (f) Once the most accurate realignment of the region is found it is reinserted into the input alignment replacing the section that was removed.

We then compare the advisors choice with the original alignment of this region, if the accuracy of the new alignment is higher we remove the columns covered by the alignment region from the input alignment and replace them with the new alignment of this region (see Figure 4f).

As a final step we compare the accuracy of the new alignment, with the realignments in the alignment regions, to the input alignment. The more accurate global alignment is returned.

### 3.3 Iterative local realignment

Once we have a new global alignment of the input sequences we can repeat the process to continue to refine the alignment. Using the same methods described earlier we compute the `Facet` score on windows of this new alignment, combine these to get column scores, and define alignment regions for which we will use parameter advising to realign. We iterate this process until a user defined maximum number of iterations is reached. Note that the local advising procedure may reach a point where none of the misaligned regions are replaced, even though continuing iteration will not effect the output we stop iterating when this happens to reduce the running time.

### 3.4 Combining local with global advising

Local advising is a method for improving the accuracy of an existing alignment. It has been shown previously that using parameter choices other than the default can greatly increase the alignment accuracy for some inputs. We can use global advising to find a more accurate starting point for adaptive local realignment.

Local and global advising can be combined in two ways.

1. **Local advising on *all* global alignments:** using adaptive local realignment on each of the alternate alignments produced within global parameter advising then choosing among all 2|*P*| alternate alignments (|*P*| unaltered global alignments and |*P*| locally advised alignments), and
2. **Local advising on *best* global alignment:** choosing the best global alignment then using adaptive local realignment to boost it’s accuracy.

We will compare both methods for combining local and global advising as well as local advising on the default alignment in the next section.

## 4 Assessing local realignment

We evaluate the performance of adaptive local realignment and its use in combination with global advising through experiments on a collection of protein multiple sequence alignment benchmarks. A full description of the benchmarks and universe of parameters used for parameter advising can be found in Kececioglu and DeBlasio (2013) and is briefly described here.

The benchmark suites used in our experiments consist of reference alignments of proteins that are largely induced by structurally aligning their known three-dimensional structure. In particular, we use the `BENCH` suite of Edgar (2009) (which is a combination of the `BAliBASE` (Bahr *et al.*, 2001), `PREFAB` (Edgar, 2004b), `OxBENCH` (Raghava *et al.*, 2003), and `SABRE` (Van Walle *et al.*, 2005) databases), supplemented by a selection from the `PALI` suite of Balaji *et al.* (2001). The full benchmark collection we use consists of 861 reference alignments.

As is common in benchmark suites, easy-to-align benchmarks are highly over-represented in this collection. To correct for this bias towards easy to align benchmarks when evaluating average advising accuracy, we binned the 861 benchmarks by *hardness*, which we measured by the true accuracy of the alignment of the benchmark’s sequences using the multiple alignment tool `Opal` under the optimal default parameter setting. We then divided the the full range [0,1] of accuracies into 10 bins, where bin b for *b* = 1,…, 10 contains hardness interval ((*b* − 1)/10, *b*/10], and has 12, 12, 20, 34, 26, 50, 62, 74, 137, and 434 benchmarks respectively. We report the average accuracy across *bins* rather than across benchmarks. This means that the average accuracy of alignments using the `Opal` default parameter settings is near 50%. Even though the binning is based on the `Opal` default alignments, most other standard aligners have default accuracy near 50%: `Clustal Omega` (Sievers *et al.*, 2011, 47.3%), `MUSCLE` (Edgar, 2004a, 48.4%), `MAFFT` (Katoh *et al.*, 2005, 51.0%). This is not to say for instance that `MAFFT` is necessarily more accurate than `Clustal Omega`, if you bin based on any aligner other than `Opal` you would get a completely new ordering. DeBlasio and Kececioglu (2015b) shows that for the task of parameter advising many of the top aligners performance nearly equally well and our choice to use `Opal` is made based on the fact that it has the highest advising accuracy in our tests. The methodology presented here is general and can be implemented for any other aligner.

**Figure 5.**
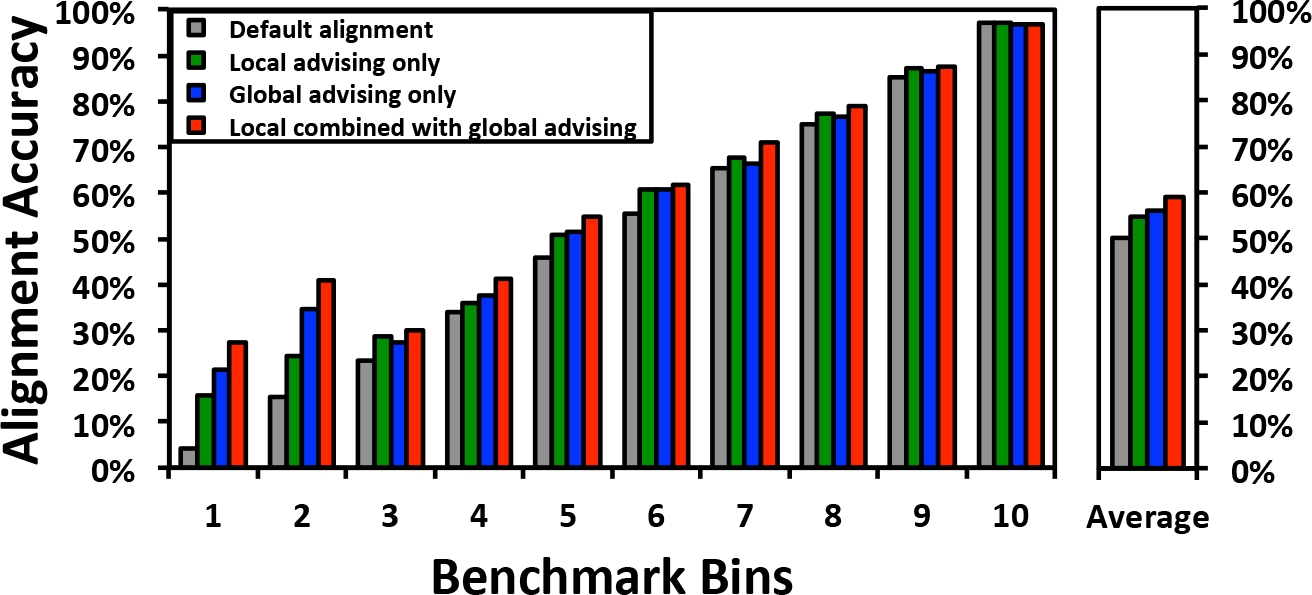
Accuracy of the default alignment, local realignment of the default alignment, parameter advising, and parameter advising with local realignment within difficulty bins. In the bar chart on the left the horizontal axis shows all ten benchmarks bins, and the vertical bars show the accuracy averaged over just the benchmarks in that bin. The accuracy of the default alignment and parameter advising using an oracle set of cardinality *k* = 10, before local realignment is shown as well as the application of local realignment to both results. The car chart on the right shows the accuracy uniformly averaged over the bins.

**Figure 6.**
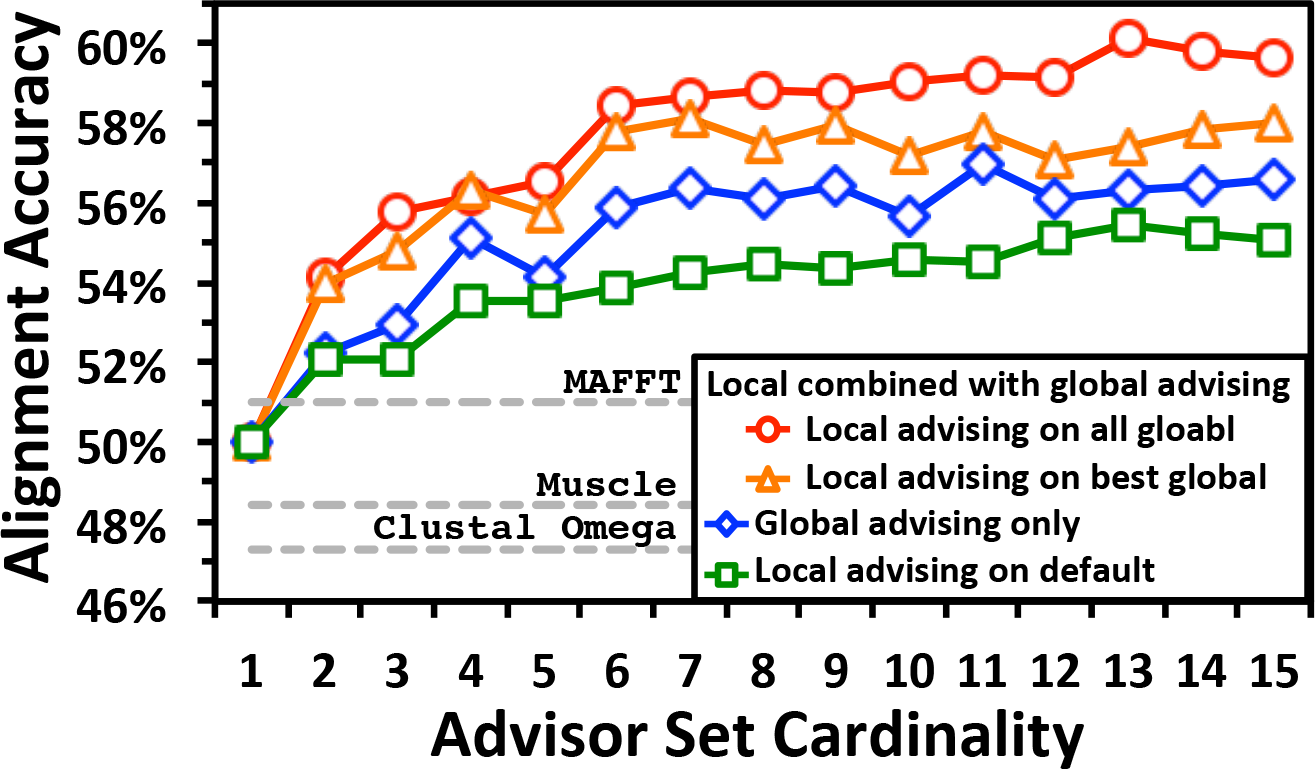
Advising accuracy using various methods versus set cardinality. This figure compares the accuracy of alignments produced by local advising on the alignment produced using the `Opal` default parameter settings, global advising alone, and two variants on combining local and global advising. The horizontal axis represents and increasing oracle advising set cardinality used for both parameter advising and local realignment. The vertical axis shows the accuracy of the alignments produced by each of the advising methods averaged across difficulty bins.

We developed a universe of alignment parameter settings U by enumerating the tunable alignment parameters within the `Opal` aligner and enumerated values from within the reasonable range of those parameters. In particular the tunable parameters for `Opal` are represented as a 5-tuple (*σ*, λ_*I*_, λ_*T*_, γ_*I*_, γ_*T*_) which represent the replacement matrix (σ) as well as the the internal and terminal gap open (λ) and extension costs (γ). For the substitution matrix we selected 3 matrices form the BLOSUM (Henikoff and Henikoff, 1992) and `VTML` (Müller *et al.*, 2002) families, three choices each for the internal and external gap open costs, three choices of internal gap extension cost, and two choices of terminal gap extension costs. We then took the cross product of the choices for each of the parameters to generate a universe of 162 parameter settings.

We use *12-fold cross validation* to examine the increase in accuracy gained using local advising both with and without the addition of global advising. We construct training and testing subsets of the alignment benchmarks by evenly and randomly distributed benchmarks into twelve groups for each hardness bin; we then formed twelve splits of the entire collection of benchmarks into a training class and a testing class, where each split placed one group in a bin into the training class and the other eleven groups in the bin into the training class; finally, for each split we generated a *training set* and *testing set* of examples alignments as follows: for each bencHmark *B* in a training or testing class, we generate |*U*| example alignments in the respective training or testing set by running `Opal` on *B* with each parameter in *U*. An estimator learned on the examples from a training set was evaluated on examples from the corresponding testing set. The results we reported are averages over twelve folds, where each *fold* is one of these pairs of associated training and testing sets. (Note that across twelve folds, every example is tested on exactly once.)

We trained the estimator coefficients for `Facet` using the difference fitting method described in Section 2.2 on the training sets described above. We found that there was very little change in coefficients between the training folds so for ease of experimentation we use the estimator coefficients that are release with the newest version `Facet` which were trained on all available benchmarks.

We examined several settings for the tunable parameters of the local realignment method: estimator window size percentage *w* (10%,20%,30%,…,90%), minimum window sizes *w_min_* (5,10, 20, 30), minimum window sizes *w_max_* (30,50,75,100,125), good and bad column label percentages *B_G_*,*B_B_* (5%,10%,20%,30%,…,70%), and gamma decay value *d* (0.5,0.66,0.9,0.99). We used the performance on *training* benchmarks described above to find the combination of these settings that gave the highest improvement in accuracy when local advising was applied to the default alignments from `Opal`. We found that using *w* = 30%, *w_min_* = 10, *w_max_* = 30, *B_G_* = 10%, *B_B_* = 30%, and d = 0.9 provided the highest increase for the training benchmarks and these are the settings we use through out the experimental results. We also iterate the local advising step five times and use this for all experiments other than the comparison to `TCS` in Section 4.3, full details why we used five iterations are shown in Section 4.4.

### 4.1 Effect of local realignment across difficulty bins

Figure 5 shows the alignment accuracy across difficulty bins for default alignments from `Opal`, local advising on these default alignments, global advising alone, and local combined with global alignment. Here the combination method uses local advising on all alternate alignments within global advising. The oracle set of cardinality *k* = 10 was used for both global and local advising.

Local advising greatly improves the alignment accuracy of default alignments (left two bars in each group). In the two most difficult benchmark bins (to the left of the figure) using local advising increases the average accuracy by 11.5% and 9.1% respectively, but the accuracy increases on all bins. Overall using local advising increases the accuracy of the default alignments by and average of 4.5% across bins.

Combining local and global advising greatly improves the accuracy over either of the methods individually. This is most pronounced for the hardest to align benchmarks. For the bottom two bins using both parameter advising and adaptive local realignment increases the accuracy by 23.0% and 25.6% accuracy over using just the default parameter choices. Additionally, using adaptive local realignment increases the accuracy by 5.9% and 6.4% accuracy on the bottom most bins over using parameter advising alone. On average thats an 8.9% increase in accuracy over all bins by using the combined procedure over using just the default parameter choice and a 3.1% increase over using only parameter advising.

### 4.2 Varying advising set cardinality

In previous sections we focused on using advising sets of cardinality *k* = 10. Because an alignment is produced for each region of local realignment for each parameter choice in the advising set it may be desirable to use a smaller set to reduce the running time of local (or global) advising. We produced oracle advising sets for cardinalities *k* = 2…15 and used them to test the effect of local advising both alone and in combination with global advising. Figure 6 shows the average advising accuracies of using advising sets of increasing cardinalities under the 3 conditions described above as well as the combination of local with global advising where the local advising step is only performed on the single best alignment identified by global advising. The cardinality of the set used for both parameter advising and local realignment is shown on the horizontal axis, while the vertical axis shows the alignment accuracy of the produced alignments averaged first within difficulty bins then across bins.

The accuracy of alignments produced by all four methods shown eventually reaches a plateau where adding additional parameters to the advising set no longer increases the alignment accuracy. This plateau is reached at cardinality *k* = 10 when local realignment is applied to the default alignments and at *k* = 6 for parameter advising with and without local realignment, but this plateau is higher for the combined methods.

Across all cardinalities using local combined with global advising improves alignment accuracy by nearly 4% on average.

The results above give average advising accuracy uniformly weighted across *bins*. We now report average advising accuracy uniformly weighted across *benchmarks*. Using its default parameter choice the `Opal` aligner achieves accuracy 80.4%. Applying both local and global advising at cardinality *k* = 10, this increases to 83.1% (performing local advising on all global alternate alignments). Using only local or global advising achieves accuracy 82.1% or 81.8% respectively. At *k* = 5 the accuracy of using local and global advising is 82.7%. By comparison, the average accuracy of other standard aligners on these benchmarks is: `Clustal Omega`, 77.3%; `MUSCLE`, 78.09%; `MAFFT`, 79.38%.

### 4.3 Comparing estimators for local advising

We have shown previously that the `Facet` estimator has the best performance for the task of global advising compared to the other accuracy estimators available (see Kececioglu and DeBlasio, 2013; DeBlasio and Kececioglu, 2015b). Figure 7 shows the average accuracy of local advising on default alignments using both `Facet` and `TCS` (the next best estimator for advising) using advisor sets of cardinality *k* = 2… 15. We found that using `TCS` for local advising greatly increased the running time because it is an additional system call and additional file IO. Because of the additional computational requirements we did not iterate the local advising for either estimator. Using `TCS` for local advising gives an increase in accuracy of less than half that of `Facet`.

**Figure 7.**
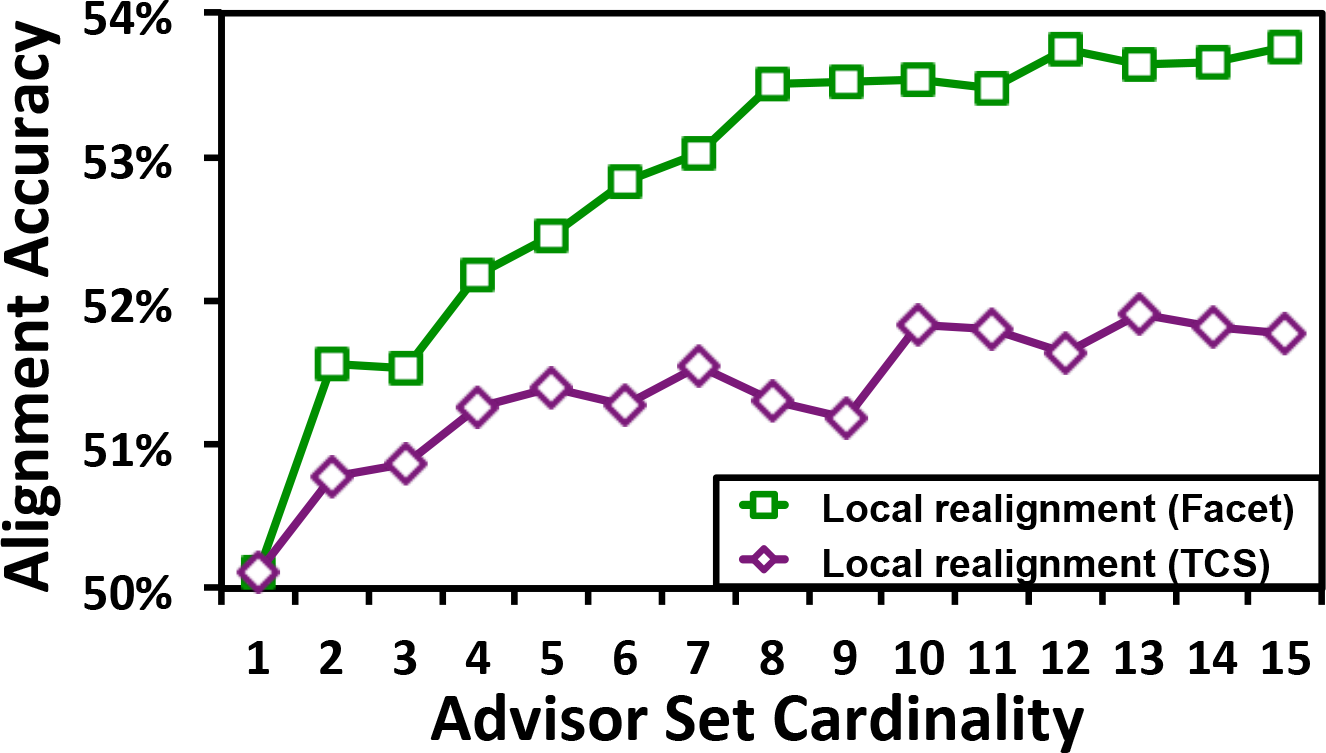
Accuracy of the default alignment and local realignment using `TCS` and `Facet` with various advisor set cardinalities. This figure compares the accuracy of alignments produced by the `Opal` default parameter settings applying local realignment using either the `TCS` or `Facet` estimator. The horizontal axis represents and increasing oracle advising set cardinality used for local realignment. The vertical axis shows the accuracy of the alignments produced by each of the advising methods averaged across difficulty bins.

### 4.4 Effect of iterating local realignment

The local advising process can be considered a refinement step for multiple sequence alignment. To continue refining the alignment we can iterate the local advising procedure (see Section 3.3). Iterating local advising should eventually reach some convergence in the alignment where you’re no longer improving the result (i.e. the alignment regions are not being changed) or even worse you start deteriorating the result due to some noise in the accuracy estimator. To find the optimal iteration limit we ran all iteration limits from 1 to 25. We found that the peak accuracy on training benchmarks was at 5 iterations, and we use that for all other experiments shown in this chapter (other than those in Section 4.3). The table below shows the average accuracy of using local adaptive realignment on the default alignment with various number of iterations.

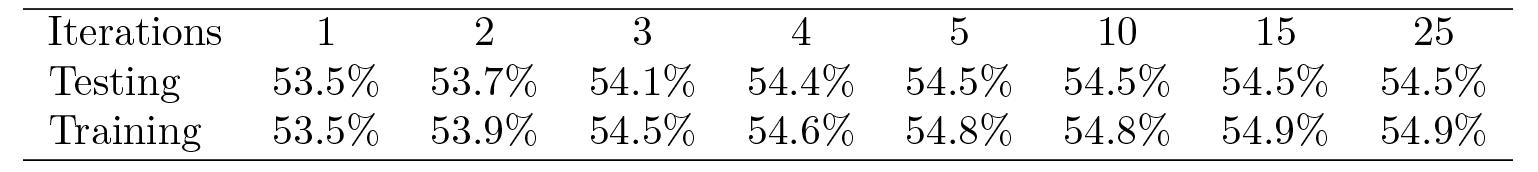

| Iterations | 1 | 2 | 3 | 4 | 5 | 10 | 15 | 25 |
| --- | --- | --- | --- | --- | --- | --- | --- | --- |
| Testing | 53.5% | 53.7% | 54.1% | 54.4% | 54.5% | 54.5% | 54.5% | 54.5% |
| Training | 53.5% | 53.9% | 54.5% | 54.6% | 54.8% | 54.8% | 54.9% | 54.9% |

### 4.5 Summarizing the effect of adaptive local realignment

Table 1 summarizes how adaptive local realignment behaves across difficulty bins when used to modify alignments produced using the default parameter setting in `Opal`. The first two rows show how many of the 861 benchmarks are in each bin, as well as how many of them had at least one realignment region where the advisor chose to replace the global alignment. The fourth row shows the average number of *Bad* in a benchmark; on average about 2 regions were realigned for each default alignment. The last two rows summarize the percentage of the original columns those *Bad* regions covered, and how many of the columns from the original alignment ended up being replaced. In the easiest-to-align benchmark bin only 47% of the alignment columns were altered, while in the rest of the bins over 60% of the alignment columns were improved.

**Table 1.** Summary of Adaptive Local Realignment on Default Alignments

| Bin | 1 | 2 | 3 | 4 | 5 | 6 | 7 | 8 | 9 | 10 | Overall |
| --- | --- | --- | --- | --- | --- | --- | --- | --- | --- | --- | --- |
| Total number of benchmarks | 12 | 12 | 20 | 34 | 26 | 50 | 61 | 74 | 137 | 434 | 861 |
| Number of benchmarks modified by adaptive local realignment | 8 | 7 | 16 | 27 | 19 | 34 | 46 | 61 | 115 | 352 | 685 |
| Percentage of benchmarks altered | 67% | 58% | 80% | 79% | 73% | 68% | 74% | 82% | 84% | 81% | 80% |
| Average number of realignment regions per benchmark | 1.92 | 2.17 | 2.50 | 1.88 | 2.23 | 2.14 | 2.31 | 2.16 | 2.48 | 2.19 | 2.23 |
| Average percentage of original columns realigned | 75% | 73% | 76% | 70% | 75% | 77% | 74% | 73% | 75% | 72% | 73% |
| Average percentage of original columns replaced | 64% | 60% | 68% | 60% | 66% | 72% | 65% | 63% | 64% | 47% | 57% |

### 4.6 Running time

As currently implemented in `Opal` local advising does not take advantage of the independence of the calls to the aligner in the parameter advising step and running them in parallel. Therefore we see a large increase in time consumption when generating locally advised alignments. In particular the average time for computing an alignment using the default global parameter setting goes from about 8 seconds to just over 36 seconds using an advisor set cardinality of *k* = 10. When iterating the local advising step 5 times we see the average running time increase to 110 seconds.

In contrast *global* advising exploits the independence of the aligner on different parameter settings. The running time for advisor set cardinality k = 10 for global advising alone is around 33 seconds, much less than the 10-fold increase to be expected if advising was not done in parallel. Even though global advising is done in parallel, local advising is not; the average running time over all benchmarks increases to 68 and 178 seconds for combining local and global advising, performing local advising on all global alignments with and without iteration, respectively.

## 5 Conclusion

We have presented *adaptive local realignment*, to our knowledge the first method that demonstrably boosts protein multiple sequence alignment accuracy by locally realigning regions that may have distinct mutation rates using different aligner parameter settings. Applying this new method alone to alignments initially computed using a single optimal default parameter setting already improves alignment accuracy significantly. When combined with methods to select an initial non-default parameter setting for the particular input sequences through global parameter advising, this new local parameter advising method greatly improves accuracy even further. We have also made available a tool that performs adaptive local realignment with the `Opal` aligner.

### Further research

For global parameter advising, *greedy advisor sets*, which are designed to work well with a given accuracy estimator, perform better than estimator-independent oracle sets (DeBlasio and Kececioglu, 2015b). For local parameter advising, however, these greedy sets found for global advising performed worse than oracle sets (which is why we used oracle sets in this study). Greedy sets may have underperformed due to the smaller universe of parameter settings explored here, or perhaps the known tendency of greedy global advising sets to not generalize as well is exacerbated when they are applied to local advising. We plan to investigate this further, by exploring a much larger universe of parameter settings, and reconsidering greedy set finding to now learn local advising sets specifically for the process of adaptive local realignment. Combining local parameter advising with *aligner advising*, which takes a set of aligners and selects both an aligner and its parameter setting, effectively performing local ensemble alignment (DeBlasio and Kececioglu, 2015a), also seems promising.

